# An evaluation of methods correcting for cell type heterogeneity in DNA methylation studies

**DOI:** 10.1101/032185

**Authors:** Kevin McGregor, Sasha Bernatsky, Ines Colmegna, Marie Hudson, Tomi Pastinen, Aurélie Labbe, Celia Greenwood

## Abstract

**Background:** Many different methods exist to adjust for variability in cell-type mixture proportions when analysing DNA methylation studies. Here we present the result of an extensive simulation study, built on cell-separated DNA methylation profiles from Illumina Infinium 450K methylation data, to compare the performance of 8 methods including the most commonly-used approaches.

**Results:** We designed a rich multi-layered simulation containing a set of probes with true associations with either binary or continuous phenotypes, confounding by cell type, variability in means and standard deviations for population parameters, additional variability at the level of an individual cell-type-specific sample, and variability in the mixture proportions across samples. Performance varied quite substantially across methods and simulations. In particular, the false discovery rates (FDR) were sometimes unrealistically high, indicating limited ability to discriminate the true signals from those appearing significant through confounding. Methods that filtered probes had consequently poor power. QQ-plots of p-values across all tested probes showed that adjustments did not always improve the distribution. The same methods were used to examine associations between smoking and methylation data from a case-control study of colorectal cancer.

**Conclusions:** We recommend surrogate variable analysis for cell-type mixture adjustment since performance was stable under all our simulated scenarios.

## Background

DNA methylation is an important epigenetic factor that modulates gene expression through the inhibition of transcriptional proteins binding to DNA [1]. Examining the associations between methylation and phenotypes, either at a few loci or epigenome-wide (i.e. the Epigenome-Wide Association Study or EWAS [2]) is an increasingly popular study design, since such studies can improve understanding of how the genome influences phenotypes and diseases. However, unlike genetic association studies, where the randomness of Mendelian transmission patterns from parents to children enables some inference of causality for associated variants, results from EWAS studies can be more difficult to interpret.

The choices of tissue for analysis and time of sampling are crucial, since methylation levels vary substantially across tissues and time. Methylation plays a large role in cellular differentiation, especially in regulatory regions [3, 4], and methylation patterns are largely responsible for determining cell-type-specific functioning, despite the fact that all cells contain the same genetic code [5].

Ideally, methylation would be measured in tissues and cells of most relevance to the phenotype of interest, but in practice such tissues may be impossible to obtain in human studies. Many accessible tissues for DNA methylation studies, such as saliva, whole blood, placenta, adipose, tumours, or many others, will contain mixtures of different cell types, albeit to varying degrees. Hence the measured methylation levels represent weighted averages of cell-type-specific methylation levels, with weights corresponding to the proportion of the different cell types in a sample. However, cell-type proportions can vary across individuals, and can be associated with diseases or phenotypes [6]. For example, individuals with auto-immune disease are likely to have very different proportions of autoimmune cells in their blood than non-diseased individuals [7, 8, 9, 10, 11], synovial and cartilage cell proportions differ between rheumatoid arthritis patients and controls [12], and associations with age have been consistently reported [13]. Hence variable cell-type-mixture proportions can confound relationships between locus-specific methylation levels and phenotypes, since these proportions are associated both with phenotype and with methylation levels [14].

In situations potentially subject to confounding, although less biased estimates of association can be obtained by incorporating the confounding variable as a covariate, this is not a perfect solution, since it may not be possible to distinguish lineage differences [14] or to accurately estimate the proportions of each cell type in a tissue sample [15, 16]. Initial studies of associations between DNAmethylation and phenotypes had largely ignored this potential confounding factor, which may have led to biased estimates of association and failure to replicate findings [17, 18].

However, in parallel with the increasing prevalence of high-dimensional methylation studies, a number of methods that can account for this potential confounding of methylation-phenotype associations have been developed or adapted from other contexts. Among those developed specifically for methylation data (Ref-based [19], Ref-free [20], CellCDecon [21], EWASher [22]), the first two were proposed by the same author (Houseman), but the first of these requires an external reference data set. Other methods were proposed in more general contexts where confounding does not necessarily result from cell-type mixtures yet is still of concern; many of these rely on some implementation of matrix decompositions (SVA [23] ISVA [24], Deconfounding [25], RUV [26, 27]).

Although there are numerous similarities between the approaches, there remain some fundamental differences in terms of limitations and performance. Unbiased comparison of methods has been difficult since true cell-type mixture proportions are unknown, replications using alternative technologies such as targeted pyrosequencing do not lead to genome-wide data where cell type proportions can be estimated, and new methods have tended to be compared with only a few other approaches. Since the problem of confounding plagues all researchers in this field, a careful comparison of existing methods correcting for cell-type heterogeneity is essential, and this is the objective of our paper.

In an ongoing study of incident treatment-naive patients with one of four systemic auto-immune rheumatic diseases (SARDs), whole blood samples were taken at presentation, and immune cell populations (purity > 95%) were sorted from peripheral mononuclear cells (PBMCs) of these patients. Analysis of DNA methylation was then performed with the Illumina Infinium HumanMethylation450 BeadChip (450K) on the cell-separated data. These data provide a unique and valuable opportunity to compare the performance of methods for cell-type mixture adjustments. We present here the results of an extensive simulation study where we remixed the cell-separated methylation profiles to incorporate variable mixture proportions and confounding of associations, and we then compare performance of 8 different methods of adjustment. We also compare the ease of use of each method, and provide an R script allowing for easy implementation of several of the best-performing methods. As far as we are aware, this is the first study to compare such an extensive set of methods in a simulation based on cell-separated data.

## Results

### Patients and original methylation profiles

Whole blood samples were obtained from patients with incident, treatment-naive rheumatoid arthritis (*n* = 11), systemic lupus erythematosus (*n* = 9), systemic sclerosis (*n* = 14), and idiopathic inflammatory myositis (*n* = 3). Several control samples were available as well (*n* = 9). CD4, CD19 and CD14 subpopulations were sorted from PBMC by magnetic cell isolation and cell separation (MACS) sorting (see Methods). Purity of the isolated populations was confirmed by flow cytometry. Only in samples with a purity > 95% methylation profiles were assessed by using the Illumina Infinium HumanMethylation 450 BeadChip on the separate cell populations. Our simulation and results are based primarily on 46 patients for whom cell sorted methylation profiles were available for both CD4+ T lymphocytes and CD14+ monocytes.

The heatmap in Figure 1 shows some representative patterns of methylation in the SARDs samples across CD14+ monocytes, CD4+ T-cells and CD19+ B-cells (this latter cell type was not available for all patients), at 200 CpG sites that were selected because of the inter-cell-type differences. The figure demonstrates that there are sizeable differences in methylation levels between cell types, and it follows that small variations in the proportions of these component cell types in a mixed tissue sample can lead to great difficulties in interpreting any phenotype-associated results.

**Figure 1:**
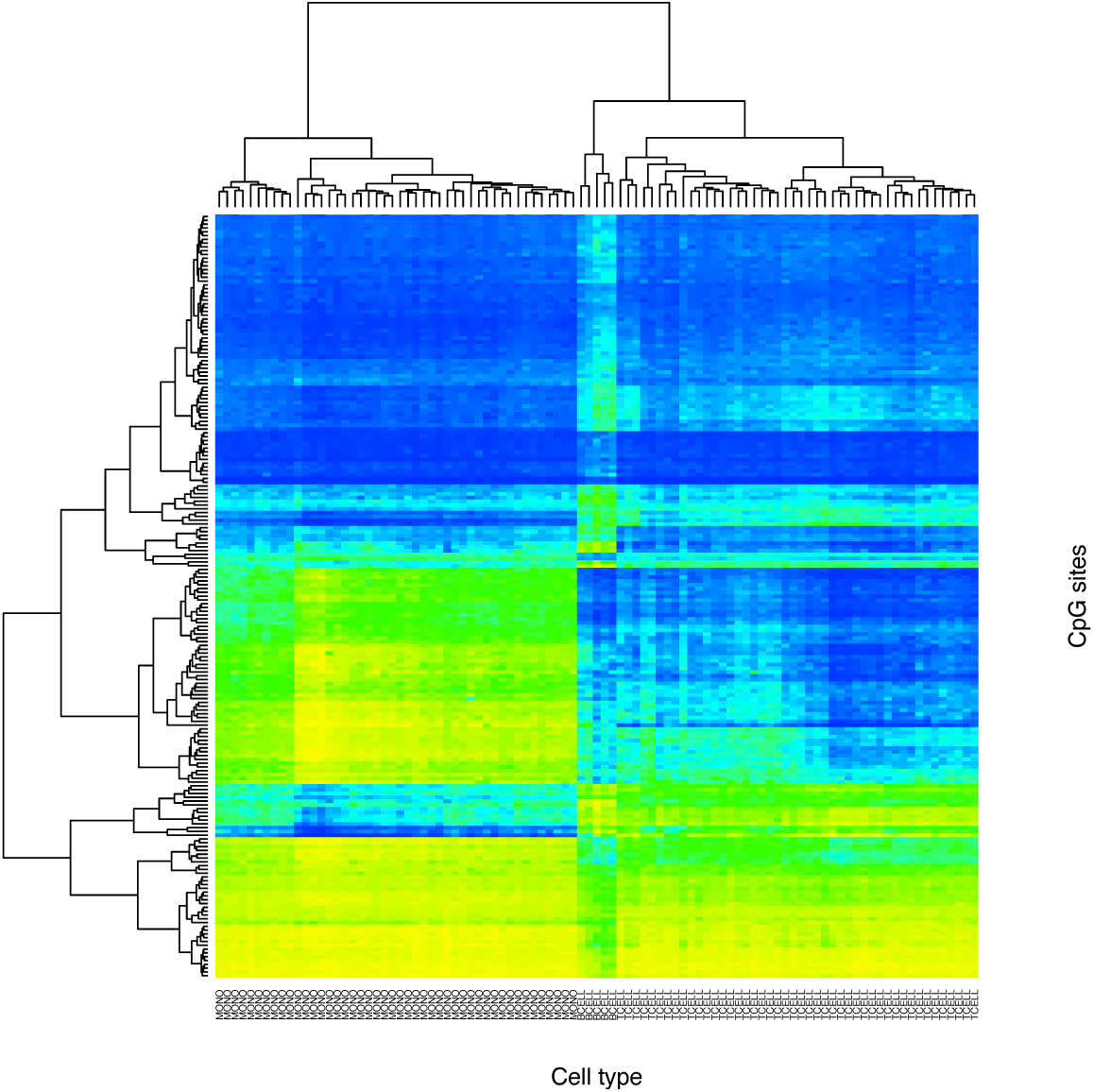
Clustered heatmap showing patterns of methylation in 46 SARDs samples (columns) and 200 CpG sites (rows), where the sites were selected to highlight the methylation differences between cell types. Consequently, the samples cluster by cell type: Monocytes, B-cells, then T-cells

### Multilayer simulation design

We implemented a rich simulation design, based on the SARDs methylation data. This simulation contains random sources of variability at multiple levels, including both variability of population mean parameters as well as variability at individual level parameters. Starting with the observed cell-separated methylation profiles for T cells and monocytes from the SARDs data, we simulated a number of probes to have associations directly with the phenotype, and we induced confounding by combining the two cell types in proportions that vary across individuals. Although this simulation design is complex and depends on a large number of parameters, it allows substantial flexibility in specifying consistency or variability between cell types, individuals or probes, and easily allows us to create realistic and pathological situations in the same framework.

Let *i* = 1, … *n*, where *n* = 46 denote the individuals in the SARDs data. In brief, the simulation proceeds as follows (see Methods for more details):

1. Select a set of 500 CpG sites where “true” associations with a phenotype will be generated by our simulation; we refer to these CpG’s as Differentially Methylated Sites (DMS).
2. Generate a phenotype, either binary (Disease/No disease) or continuous.
3. For any probe not in the DMS set, the cell-type-specific methylation values are the observed values from the real data.
4. For a probe in the DMS set, a randomly-generated quantity is added to the observed cell-type-specific level of methylation, in a way that depends on the phenotype.
5. The cell-type-specific methylation values are mixed together in proportions that vary depending on the phenotypes

Over all DMS, one would expect to see a range of positive and negative associations with the phenotype. In step 4, we allow these associations to differ between cell types in order to specify an association between change in methylation and cell type. After having specified each of the site and cell type specific associations, we then add between-subject variability to each site. The final step, #5, leads to methylation proportions as they would appear had the mixed tissue been analyzed directly.

After simulating data, we then test for association between the phenotype and the methylation levels in the mixed data at each probe. We compare the p-values obtained from these tests of association across 6 simulation scenarios and 8 different methods for cell type adjustment. Some of the DMS were simulated to have very small effects and therefore statistical tests of association may not be significant. On the other hand, since the cell-type proportions vary with phenotype, this can lead to non-DMS sites showing spurious associations with the phenotype. Notation for key parameters is given in Table 1, and the parameter choices across different simulation scenarios are summarized in Table 2.

**Table 1:** Fixed parameters in the simulation design

| Parameter | Description |
| --- | --- |
| $S$ | Number of CpGs chosen to be associated with phenotype in simulation |
| $\mu_k$ | Mean of the cell-type-specific DMS effects for cell type $k$, over all $S$ DMS |
| $\sigma_k$ | Standard deviation of cell-type-specific DMS effects for cell type $k$, over all $S$ DMS |
| $\sigma_{jk}$ | Variability of individual deviations |
| $\alpha^{(0)} = (\alpha_1^{(0)}, \alpha_2^{(0)})^\top$ | Expected proportion of the mixture from cell types 1 and 2 when the phenotype $z_i$ is zero. |
| $\alpha^{(Z)} = (\alpha_1^{(Z)}, \alpha_2^{(Z)})^\top$ | Average cell type mixture proportions for cell types 1 and 2 for subjects with phenotype level $Z$ (continuous or binary) |
| $\rho$ | Precision of simulated cell mixture distributions. A greater value corresponds to more clearly defined differences in cell type proportions with respect to the phenotype |

**Table 2:**
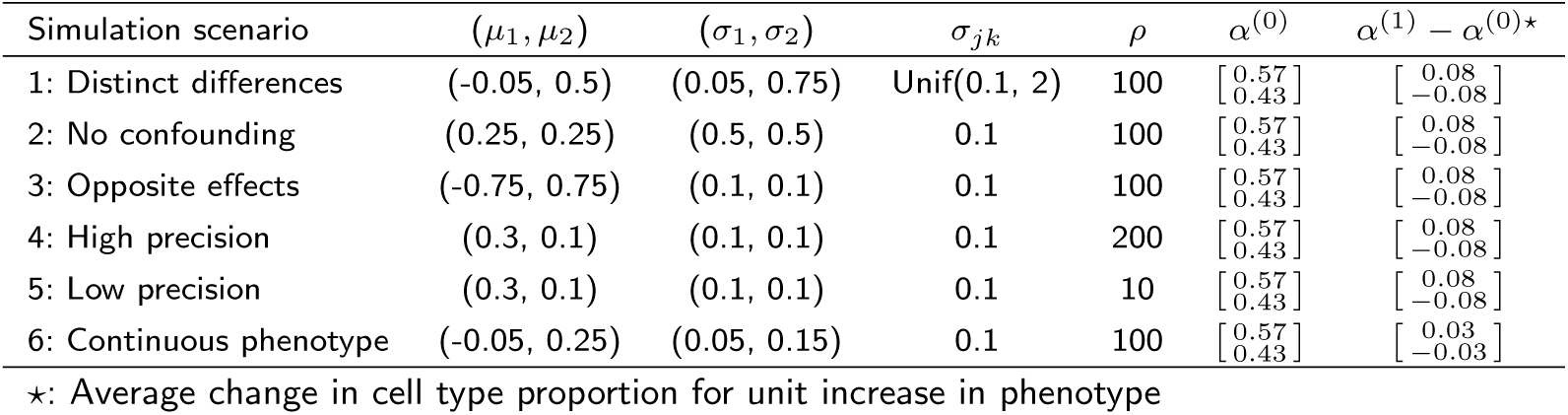
Parameter choices for the simulation scenarios. *k* = 1 corresponds to monocytes, and *k* = 2 corresponds to CD4+ T-cells.

### Scenario 1: DMS sites have differences in both means and variances between cell types

In our first simulation scenario (Table 2), we chose to specify distinct differences in the strength and distribution of the methylation-phenotype associations (DMS) for the two cell types, with a binary phenotype. Differences in the DMS distributions include both direction of the effects as well as the amount of variability across sites and individuals; Supplement Figure 1 displays histograms of the 500 simulated values of the DMS means *μ_jk_* for the two cell types, showing the substantial differences between these two distributions.

In this scenario, we would expect that an analysis not taking cell type into consideration should result in many p-values that are smaller than expected, or equivalently a greatly inflated slope in a p-value QQ-plot, due to the strong confounding built into the simulation design. After performing analysis for all probes with unadjusted data, we repeated the epigenome-wide analysis with 8 popular or newly developed adjustment methods (see Materials and Methods). QQ-plots for these 8 methods as well as the uncorrected analysis can be seen in Figure 2. Examination of the left hand side of this plot (x-axis smaller than about 3.5) shows that there is, indeed, a genome-wide inflation of p-values in the analysis uncorrected for cell-type mixture. Encouragingly, most methods do a fairly good job in correcting for the confounding, since the corrected QQ plots are close to the line of expectation up until the tails of each set of p-values. The reference-free method, however, continues to display inflation even after correction.

**Figure 2:**
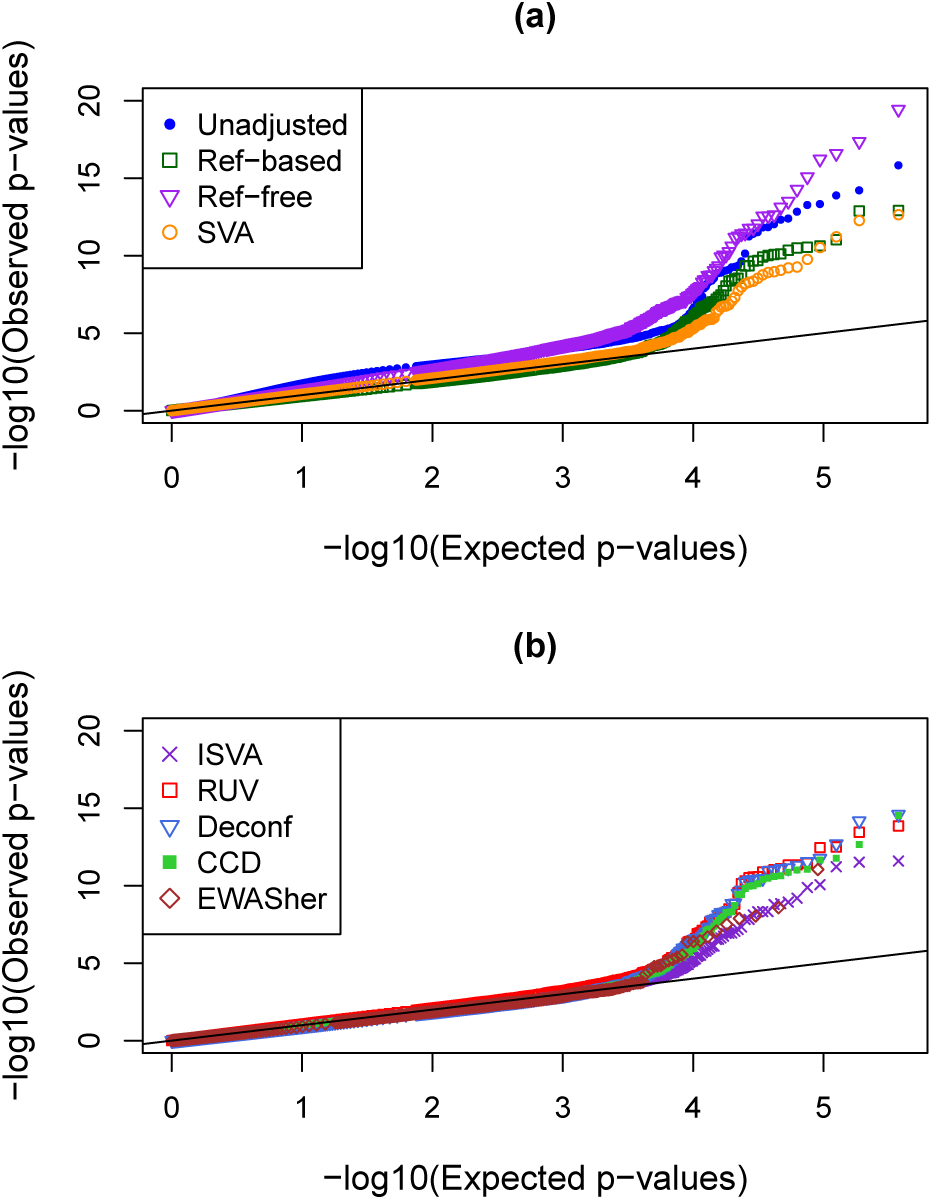
QQ-plots showing distributions of p-values in simulation Scenario 1 where the true effects in the different cell types have very distinct distributions. Results are shown with no adjustment for cell type mixture as well as with 8 other methods; these are split across two panels to clarify the display

Several numeric performance metrics can be seen in Table 4a for this simulation scenario. One of these metrics is the genomic inflation factor (GIF) [28], which is the slope of the lines seen in Figure 2 after removing the 500 DMS. The unadjusted GIF was 1.6, indicating a substantial inflation of significance across all p-values, but after adjustment most values are quite close to 1.0, as would be expected in the absence of any confounding. The Ref-based, EWASher and Deconf methods have slopes slightly less than 1.0, implying possible overcorrection.

**Table 3:**
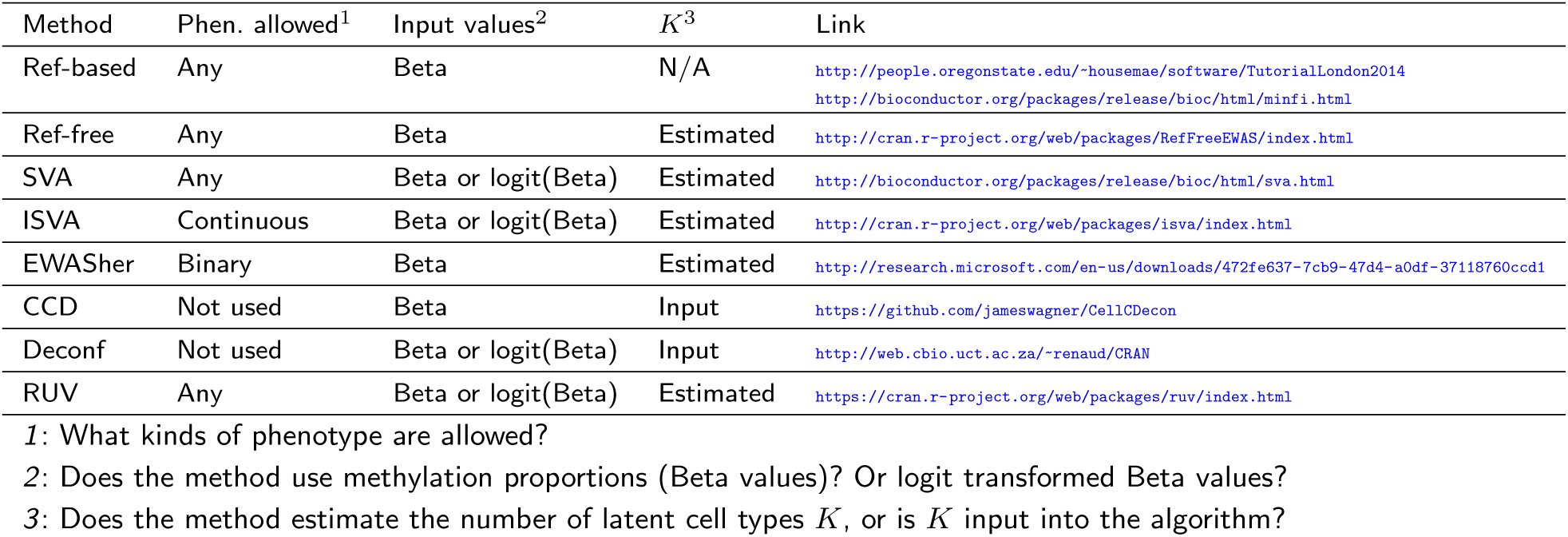
Comparison of some features of the methods for cell type mixture adjustment

**Table 4:** Performance metrics under all simulation scenarios

(a) Scenario 1. Distinct associations between cell types
| Method | FDR | Power | KS<sup>1</sup> | GIF<sup>2</sup> | $\hat{K}$<sup>3</sup> |
| --- | --- | --- | --- | --- | --- |
| Unadjusted | 0.179 | 0.096 | 0.1679 | 1.60 | - |
| Ref-based | 0.021 | 0.094 | 0.0255 | 0.92 | - |
| Ref-free | 0.53 | 0.206 | 0.0965 | 1.33 | 13 |
| SVA | 0.054 | 0.07 | 0.0205 | 1.05 | 10 |
| ISVA | 0 | 0.078 | 0.0035 | 0.97 | 12 |
| EWASher | 0.071 | 0.026 | 0.0437 | 0.92 | - |
| CellCDec | 0.021 | 0.094 | 0.0084 | 0.98 | - |
| Deconf | 0.037 | 0.104 | 0.023 | 0.93 | - |
| RUV | 0.017 | 0.116 | 0.0184 | 1.04 | 43 |

(d) Scenario 4. High precision
| Method | FDR | Power | KS | GIF | $\hat{K}$ |
| --- | --- | --- | --- | --- | --- |
| Unadjusted | 0.524 | 0.594 | 0.1423 | 1.32 | - |
| Ref-based | 0.161 | 0.406 | 0.0432 | 1.04 | - |
| Ref-free | 0.564 | 0.698 | 0.1007 | 1.36 | 13 |
| SVA | 0.136 | 0.494 | 0.0318 | 1.09 | 11 |
| ISVA | 0.297 | 0.482 | 0.0618 | 1.20 | 14 |
| EWASher | 0.118 | 0.06 | 0.0383 | 0.94 | - |
| CellCDec | 0.0781 | 0.472 | 0.0312 | 1.05 | - |
| Deconf | 0.0927 | 0.47 | 0.0138 | 1.02 | - |
| RUV | 0.265 | 0.542 | 0.0508 | 1.15 | 32 |

(b) Scenario 2. No confounding
| Method | FDR | Power | KS | GIF | $\hat{K}$ |
| --- | --- | --- | --- | --- | --- |
| Unadjusted | 0.0246 | 0.632 | 0.059 | 1.09 | - |
| Ref-based | 0.0171 | 0.572 | 0.0296 | 0.90 | - |
| Ref-free | 0.499 | 0.702 | 0.0581 | 1.25 | 14 |
| SVA | 0.120 | 0.628 | 0.0148 | 1.06 | 10 |
| ISVA | 0.551 | 0.654 | 0.0613 | 1.27 | 15 |
| EWASher | 0 | 0.122 | 0.0662 | 0.83 | - |
| CellCDec | 0.0126 | 0.626 | 0.0205 | 0.93 | - |
| Deconf | 0.0792 | 0.628 | 0.0313 | 0.92 | - |
| RUV | 0.229 | 0.624 | 0.0074 | 1.00 | 38 |

(e) Scenario 5. Low precision
| Method | FDR | Power | KS | GIF | $\hat{K}$ |
| --- | --- | --- | --- | --- | --- |
| Unadjusted | 0.141 | 0.586 | 0.241 | 1.47 | - |
| Ref-based | 0.456 | 0.636 | 0.1975 | 1.49 | - |
| Ref-free | 0.686 | 0.7 | 0.1453 | 1.48 | 12 |
| SVA | 0.201 | 0.614 | 0.0911 | 1.23 | 6 |
| ISVA | 0.0876 | 0.396 | 0.0404 | 1.12 | 11 |
| EWASher | 0 | 0.042 | 0.0299 | 0.92 | - |
| CellCDec | 0.243 | 0.552 | 0.0082 | 1.27 | - |
| Deconf | 0.249 | 0.62 | 0.2038 | 1.41 | - |
| RUV | 0.976 | 0.784 | 0.2282 | 1.93 | 33 |

(c) Scenario 3. Opposite effects
| Method | FDR | Power | KS | GIF | $\hat{K}$ |
| --- | --- | --- | --- | --- | --- |
| Unadjusted | 0 | 0.462 | 0.0408 | 0.97 | - |
| Ref-based | 0 | 0.484 | 0.0362 | 0.87 | - |
| Ref-free | 0.259 | 0.594 | 0.0531 | 1.19 | 14 |
| SVA | 0.0582 | 0.55 | 0.0034 | 1.00 | 10 |
| ISVA | 0.0521 | 0.546 | 0.0063 | 1.01 | 14 |
| EWASher | 0 | 0.108 | 0.0915 | 0.74 | - |
| CellCDec | 0.0083 | 0.48 | 0.0326 | 0.88 | - |
| Deconf | 0 | 0.492 | 0.0346 | 0.89 | - |
| RUV | 0.0194 | 0.504 | 0.0118 | 1.00 | 37 |

(f) Scenario 6. Continuous
| Method | FDR | Power | KS | GIF | $\hat{K}$ |
| --- | --- | --- | --- | --- | --- |
| Unadjusted | 0.163 | 0.514 | 0.081 | 1.37 | - |
| Ref-based | 0.010 | 0.44 | 0.031 | 0.99 | - |
| Ref-free | 0.160 | 0.6 | 0.0606 | 1.17 | 14 |
| SVA | 0.045 | 0.464 | 0.006 | 0.99 | 10 |
| ISVA | 0.166 | 0.494 | 0.076 | 1.21 | 14 |
| CellCDec | 0.590 | 0.406 | 0.079 | 1.44 | - |
| Deconf<sup>4</sup> | - | - | - | - | - |
| RUV | 0.023 | 0.518 | 0.006 | 0.966 | 39 |

<sup>1</sup> *KS*: Kolmogorov-Smirnov statistic; <sup>2</sup> *GIF*: Genomic inflation factor; <sup>3</sup> $\hat{K}$ : Estimated latent dimension; <sup>4</sup> No results were obtained with $K=3$ in the allowable time on the computational cluster *mammoth*

Due to the large number of statistical tests performed, all p-values corresponding to the CpG-phenotype associations undergo the Benjamini-Hochberg procedure to control for the false discovery rate (FDR) [29]. Adjusted p-values falling below the 0.05 threshold after this correction are considered significant. Since we know which 500 sites were generated to be truly DMS, Table 4a reports both the power—i.e. the proportion of these DMS sites that were identified with adjusted *p* < 0.05—and also the proportion of all the other sites that would be declared significant, i.e. the false discovery rate (FDR). This table also shows a measure of performance based on the Kolmogorov-Smirnov (KS) test for whether the p-value distribution matches the expected uniform distribution. Of course, the KS test assumes independence of all the individual tests, and therefore we are not using this test in order to perform inference, but simply as a measure of deviation where smaller values imply less deviation.

For this simulation (Scenario 1 with distinct association distributions in the two cell types) all methods except the reference-free method achieved an improved (smaller) FDR than the unadjusted analysis for the non-associated probes. Using ISVA, the FDR was zero, and it was under 5% for the Ref-based, CellCDec, Deconf, and RUV methods. Although the power (sensitivity) for all the methods appears very low, many of the simulated effect sizes at the chosen DMS sites were very small, and nevertheless the rankings of the different methods is still informative. Supplemental Figure 1 shows the simulated means of the cell-type distributions for the 500 probes; subsequently, additional random errors were introduced at the level of each individual leading to substantial variability in the realized methylation differences. The power of most methods was more or less on par with the unadjusted analysis, except for Ref-free, EWASher and RUV. Power for EWASher is extremely poor; this method removes probes with very high or very low levels of methylation prior to constructing the components, and hence many DMS probes are not even included in analyses. Results from the Ref-free power must be interpreted cautiously since the type 1 error is so substantially elevated for this method. The RUV method’s FDR is surprising given that the QQ-plot looks very similar to CellCDec and the other statistics in Table 4a are good; possibly a large number of the DMS probes are being captured for constructing of the deconfounding components. The KS statistic confirms the conclusions obtained from other metrics, showing small values for most methods except for the unadjusted data and the Ref-free method.

### Scenario 2: No confounding

It is also of interest to examine performance when there is no confounding. By simulating data with the same cell-type-specific means and variances in both cell types (Scenario 2, Table 2b), the unadjusted analysis should not be subject to any bias. As expected, Table 4b shows low FDR and good power for the unadjusted method, and similar results are obtained for CellCDec, Deconf and Ref-based. In Supplemental Figure 2, it can be seen that the unadjusted results lie very close to the line of expectation, apart from the tail of the distribution where the DMS predominate. It is interesting to note that SVA, RUV, Ref-free and in particular ISVA display high FDR values, implying that far too many DMS probes are being inferred. Despite the fact that no confounding was simulated, the GIF for the unadjusted data is slightly inflated; in fact, after adjustment, the GIF increases for Ref-free and ISVA. In contrast, the GIF is smaller than one for CellCdec, Deconf and the Ref-based methods, implying some over-correction.

### Scenario 3: Opposite effects in different cell types

To investigate a case of severe differential effects, the cell-type-specific means were selected to have opposite signs in the two cell types in Scenario 3. In this case, the mixed sample can have small DMS effects, since the two cell-type-specific effects may cancel each other. Confirming this expectation, there is no inflation of the test statistics in the unadjusted data (GIF=0.97). Similar to the previous scenarios, we see very poor power and overcorrection with EWASher, and extremely inflated FDR with the Ref-free method. Small power improvements over the unadjusted analysis can be seen when using any of the other methods.

### Scenarios 4 and 5: Altered precision simulations

Two scenarios were generated where we changed the precision of the individuals’ cell type distributions between cases and controls. That is, a higher precision corresponds to a more pronounced separation in the cell type distributions between cases and controls, while a lower precision makes the two distributions more difficult to distinguish. Here, both T-cells and monocytes were chosen to have distinct, positive net association with the phenotype, however the precision parameter, *ρ*, from the Dirichlet distribution was varied such that *ρ* = 200 for high precision and *ρ* =10 for low precision. QQ-plots are shown in Supplemental Figures 4 (high precision) and 5 (low precision), and numeric metrics are in Tables 4d and 4e.

In the high-precision scenario, the FDR rate is extremely high when no adjustment is used. Most methods, however, perform quite well in reducing the GIF and KS statistics, reducing the FDR and retaining decent power (with the exceptions of Ref-free and EWASher as seen previously). In contrast, for the low precision scenario, where there is much more variability from one individual to the next in the mixture proportions, as well as substantial differences between cases and controls, performance is generally poor. The QQ plots display substantial inflation, and most methods have very high FDR. Even the Ref-based method has very high FDR, and notably the QQ-plot for RUV has enormous inflation and the FDR is 97%. In fact, the unadjusted analysis appears to be one of the better choices here, with lower FDR and good power; CellCDec and Deconf also seem to have better performance than the others.

### Scenario 6: Continuous phenotype simulation results

In our simulation with continuous phenotypes, the relative performances of the methods are different again. Table 4f and Supplement Figure 6 indicate that unlike all the other scenarios, the Ref-free method performs fairly decently in this case, leading to small reductions in the GIF and KS statistics and a small improvement in power. RUV’s performance is one of the best choice here, with low FDR, good power, and an excellent GIF value. In contrast, the CellCDec method, which had performed quite well in all the other scenarios, shows extensive inflation across the QQ plots (GIF=1.44) and very high FDR estimate; we were unable to obtain results for the Deconf method with 3 components within the available computational time limits on the Mammouth Compute Canada cluster. We note that the EWASher method does not allow continuous phenotypes and cannot be used.

### Estimated latent dimension

Our simulation is based on complex mixtures of methylation profiles from two separated cell types. It is therefore interesting to note that all the methods that provide estimates for the latent dimension (the last column of all subtables in Table 4) consistently provide estimates that are much larger than 2. Estimates are obtained for the Ref-free, SVA, ISVA and RUV methods. Both SVA and ISVA assume the number of surrogate variables is less than or equal to the number of true confounders whose linear space they span, and for RUV, the authors themselves commented that the estimated values for *K* do not necessarily reflect the true dimension [27]. All estimates are generally greater than 10, and RUV’s estimates tend to be over 30. In fact, there may be some additional sources of variation present in the original cell-separated methylation data, and these factors are likely being captured by these numerous latent variables. In fact, analyses of the original cell-type-separated data using patient age as the predictor resulted in estimated latent dimensions that were themselves large. For example, random matrix theory [30] (which is used for dimension estimation in the reference-free method and ISVA) estimated a latent dimension of 10 for both T-cells and monocytes when analyzed separately. Furthermore SVA estimated the number of surrogate variables to be 7 and 9 for T-cell and monocytes, respectively.

### Results from analysis of the ARCTIC dataset

We tested the performance of these 8 adjustment methods on 450K measurements from the Assessment of Risk in Colorectal Tumors in Canada (ARCTIC) study [31], and the methylation data are deposited in dbGAP under accession number [phs000779.v1.p1]. We analyzed only 977 control subjects from this study, restricting to those where DNA methylation was measured on lymphocyte pellets, and examined the association between smoking (ever smoked) and methylation levels at all autosomal probes who passed quality control (473,864 probes). We excluded the colorectal cancer patients from this analysis due to concerns that their methylation profiles may have been affected by treatment. Patient age was included as a covariate in all analyses.

Figure 3 shows the QQ-plots for 7 adjustment methods, and Table 5 provides Kolmogorov-Smirnov and GIF numeric metrics of performance. The Deconf and CellCDec methods could not be used in these data since the computational time exceeded the 5 day limit allowed on the Mammouth cluster of Calcul Quebec. As was seen in our simulations, the EWASher method seems to overcorrect, leaving no significant probes, and the GIF is much smaller than 1.0. However, all other methods lead to QQ-plots where the slope is larger after correction than before – the GIF estimates are substantially larger than for the unadjusted analysis. For SVA and ISVA, the KS-statistic is increased after corrections are applied. Furthermore, among the top 1000 probes selected by each method (based on raw p-value), none were shared by all methods (including unadjusted results). If EWASher was excluded, 87 probes overlapped among the most significant 1000, and 89 probes overlapped among methods excluding EWASher and the unadjusted results. Therefore, the methods are highlighting quite different results for the most significant probes.

**Figure 3:**
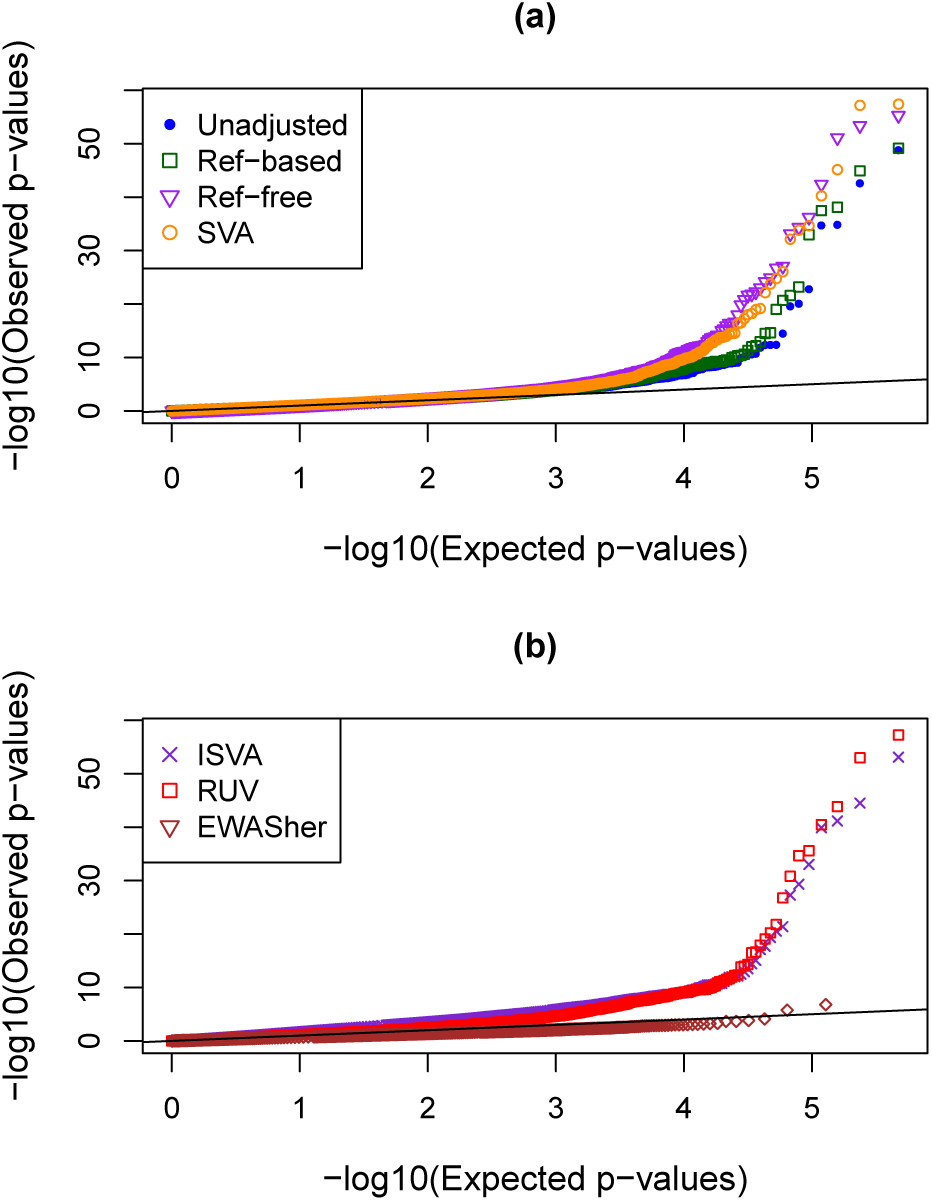
QQ-plots of — log_10_ p-values from the ARCTIC study with different adjustment methods.

**Table 5:** Performance metrics for the ARCTIC data with the most significant probes removed (top 5%). It was not possible to obtain results for the CellCDec and Deconf methods.

| Method | KS Stat | GIF | $\hat{K}$ |
| --- | --- | --- | --- |
| Unadjusted | 0.0758 | 0.9142 | - |
| Ref-based | 0.0508 | 0.9907 | - |
| Ref-free | 0.0296 | 1.1499 | 32 |
| SVA | 0.0362 | 1.0962 | 15 |
| ISVA | 0.1208 | 1.6820 | 39 |
| EWASher | 0.7291 | 1.0164 | - |
| RUV - 3 components | 0.0906 | 1.2423 | - |

In Table 6, *p*-values are shown for 7 probes that have been reliably associated with smoking [32]. Although substantial evidence of association can be seen at all probes, it is interesting to see the differences in significance across methods. For example, at probe *cg21161138*, significance ranges from 10^−7^ to 10^−25^.

**Table 6:** P-values for sites previously found to be associated with smoking. [32]

| Site | Unadj | Ref-Based | Ref-free | SVA | ISVA | EWASher<sup>1</sup> | RUV |
| --- | --- | --- | --- | --- | --- | --- | --- |
| cg06644428 | 2.83E-20 | 2.04E-21 | 1.96E-27 | 9.19E-27 | 4.41E-20 | - | 1.64E-22 |
| cg05951221 | 2.72E-43 | 1.21E-45 | 5.41E-56 | 3.96E-58 | 3.20E-45 | 5.77E-01 | 1.05E-53 |
| cg21566642 | 1.72E-49 | 6.93E-50 | 4.20E-54 | 6.59E-58 | 8.09E-54 | 6.45E-01 | 5.94E-58 |
| cg01940273 | 2.09E-35 | 3.30E-38 | 4.07E-43 | 7.07E-46 | 6.93E-42 | 3.07E-01 | 1.45E-44 |
| cg19859270 | 4.91E-13 | 2.52E-22 | 4.76E-35 | 2.21E-35 | 5.15E-28 | - | 2.16E-35 |
| cg05575921 | 1.58E-35 | 7.79E-39 | 7.89E-52 | 5.65E-41 | 1.34E-40 | - | 3.43E-41 |
| cg21161138 | 2.43E-07 | 2.99E-10 | 1.24E-25 | 1.68E-25 | 2.84E-21 | 9.05E-01 | 6.07E-21 |
| cg06126421 | 1.78E-23 | 1.06E-33 | 7.38E-34 | 7.37E-33 | 9.69E-34 | 7.05E-01 | 2.55E-36 |
| cg03636183 | 9.27E-21 | 6.74E-24 | 6.10E-37 | 1.51E-34 | 4.77E-30 | 3.09E-01 | 1.60E-31 |

<sup>1</sup>Several sites were filtered out by EWASher

### Computational performance

To compare computational time across the different adjustment methods, we selected a random sample of 10,000 CpGs from the ARCTIC methylation matrix to create a benchmark dataset. As we are not making any statistical inference here, all samples were included, regardless of whether we had matching cell type sets or quality control status. Some of the methods calculate p-values and parameter estimates internally, and others require the use of an external function to perform a linear fit. Therefore, to make the computational times comparable, we define start time as when the adjustment method is first called, and end time when all estimates and p-values have been obtained.

Figure 4 shows the running times on the log scale, as the sample size increases (N = 50 to N = 500), and-for methods where a value of the latent dimension *K* is required-as *K* increases with a fixed sample size (N = 50). There are major differences in running times for the cell type adjustment methods. Not surprisingly, the Ref-based method is very fast, as is RUV. The slowest methods are Deconf and CellCDec; the computational time required for the Ref-free method also increases quickly with the sample size. In panel (b) of Figure 4, it is interesting to note that increasing *K* has very little effect on the speed of four of the six methods that require a specification of *K*. However, both the computational time for CellCDec and Deconf increase exponentially with larger values of *K*; as noted previously, we were not able to obtain results for these methods on the ARCTIC data set when using all autosomal probes. We note also that the complexity of preparation of the input files also varies from one algorithm to another.

**Figure 4:**
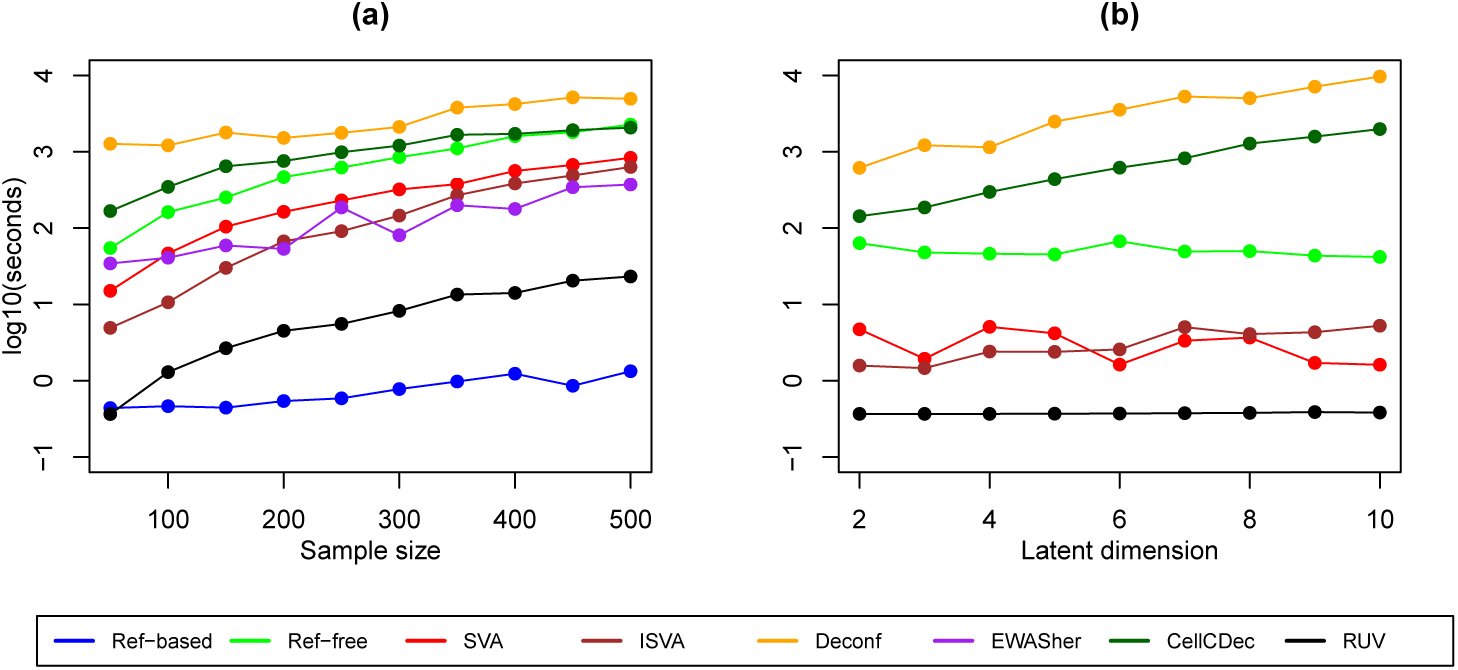
Computational time comparison by (a) sample size and (b) latent dimension. In Panel (a), the latent dimension was estimated by the algorithms as needed. In panel (b), the sample size is fixed at 50

## Discussion

We have presented an extensive comparison of 8 different methods for adjusting for cell-type-mixture confounding, by designing a rich simulation based on cell-type-separated methylation data in SARDs patients. Our simulation contained multiple levels of variability, between cell types, at the level of the probe means, and at the level of the individual. We found that there was no adjustment method whose performance was uniformly the best, and in fact in some of our scenarios, the unadjusted results were quite comparable to the best adjusted results.

In many of our simulated scenarios, as might be expected, the Ref-based method performed well. This method is very easy to implement, and as seen in the computing performance section, it runs very quickly, even on larger sample sizes. It usually achieved good statistical power, and, with one exception, reduced the FDR from the unadjusted model. It also has the advantage of being able to directly estimate the cell type composition of each sample. Therefore, the Ref-based method is an obvious choice when a complete set of the required cell-separated methylation profiles are available, however, this is not always the case. For some tissues, cell types that are of particular interest are very difficult or impossible to extract; one example here would be the syncytiotrophoblast cells in placenta [33, 34].

In every case we examined, EWASher did a very good job in reducing p-value inflation, and GIF values were substantially reduced from the unadjusted analyses. However, the fact that this method so strictly forces the GIF factor downwards may raise concerns about overcorrection. If there were, for example, global hyper-methylation associated with a disease, adjustment using EWASher would be overly conservative. Additionally, part of the algorithm involves filtering out loci that are unilaterally high or low among all subjects. The assumption behind this filter is that these loci are, for all intents and purposes, completely methylated or unmethylated and any associations between these probes and the phenotype are not interesting. This may be an overly strong assumption. In our simulations, this filter results in the dramatically worse power of this method, since we did not restrict the randomly-selected DMS loci to any particular mean level of methylation. Furthermore, the EWASher method is quite difficult to implement. Although most methods can be run in R (www.cran.r-project.org) EWASher requires the user to create three separate input files for a stand-alone executable, and then to perform post-processing in R.

We cannot explain the poor performance in our simulations of the Ref-free method. The FDR rates were almost always more inflated than in the raw data, and this inflation is clearly visible in the QQ-plots. Furthermore, implementation was somewhat more complex since the approach involved one step to estimate the latent dimension, a second to get parameter estimates, and then finally bootstrap calculations to obtain standard errors. Performance of the Ref-free method was good for Scenario 6 with a continuous phenotype, so we hypothesize that there are some linearity assumptions in the correction that are being violated in our binary phenotype simulations.

Performance of CellCDec and Deconf were generally quite good for binary phenotpes. The CellCDec method exists as a C++ program, and was quite easy to implement. The number of latent cell types must be specified in advance, which is a limitation. The run time was longer for this algorithm than the others, and increased quickly with the assumed number of cell types-in fact, we were unable to obtain results in the ARCTIC data. CellCDec does not use phenotype information; it would be interesting to see how this program would perform if it took the phenotype and other covariates into account. For Deconf, the most important limitation was the running time. In all cases, it took longer to run than the other adjustment methods, and we were unable to obtain results for the ARCTIC data. Run time was sensitive to both increases in sample size and number of cell types. Akin to CellCDecon, the fact that it does not internally estimate the number of cell types is an issue.

The results for ISVA and RUV were often among the better ones with a couple of notable exceptions: FDR rates were extremely high for RUV in the low precision scenario, and for ISVA in the no confounding scenario. Computational time for the ISVA method also increased quite rapidly with sample size. RUV is very easy to run and is available as an R function. It contains a function to estimate the latent dimension (*K*), although, akin to the other methods that estimate *K*, the estimated dimension tends to be much higher than the simulated reality. We performed some investigations into how RUV performs at a range of values for *K*, and the best performance was observed, in most simulation scenarios, at smaller values such as *K* = 3; recently Houseman has also found that estimated latent dimsions obtained through random matrix theory may not be the best choices [35]. RUV is also extremely fast-slower only than the Ref-based method, and as shown in Figure 4, the computational time is essentially invariant as the latent dimension is varied, making this an attractive option. Nevertheless, the SVA method, although rarely the best, was a method that did not have any notable failures across our scenarios, and was easy to implement.

There are other methods for deconvolution that we did not examine, especially in the computer science and engineering literature [36]. However, it is not clear whether these methods would be easily adapted for use on methylation data. Also, new meth for DNA methylation analysis continue to be published, such as [37]. However the spectrum of methods that we have examined includes the most-commonly used approaches. All methods that we have examined assume approximately linear relationships between the phenotype and the methylation levels or covariates; however, this should not be an important limitation since approximate linearity should hold [35].

The latent dimension, when estimated, was rarely similar to the dimension of *K* = 2 implemented in our simulation. However, these estimates of *K* capture aspects of heterogeneity in the data that are only partially attributable to the mixture of data from two cell types. This heterogeneity may also be partially due to technological artefacts from batch effects or experimental conditions, and in particular to the fact that more subtle cell lineage differences will still be present even after cell sorting [35].

In summary, our simulation study comparing methods found a wide range of performance across our scenarios with notable failures of some methods in some situations. We recommend SVA as a safe approach for adjustment for cell-type mixture since it performed adequately in all simulations with reasonable computation time. A set of scripts enabling implementation of all these methods can be found at https://github.com/GreenwoodLab/CellTypeAdjustment.

## Conclusions

We have compared 8 different methods for adjusting methylation data for cell-type-mixture confounding in a rich and multi-layered simulation study, and in a large set of samples where methylation was measured in whole blood. No method performs best in all simulated scenarios, nevertheless we recommend SVA as a method that performed adequately without notable failures.

## Author Contributions

Study design: KM, AL, CG; Data collection: MH, SB, IC, TP; Data analysis: KM; Writing manuscript: KM, AL, CG, MH.

## Acknowledgements

This work was funded by the Ludmer Centre for Neuroinformatics and Mental Health and by Canadian Institutes of Health Research operating grant MOP-300545. Computations were made on the supercomputer Mammouth parallèle 2 from Université de Sherbrooke, managed by Calcul Québec and Compute Canada. The operation of this supercomputer is funded by the Canada Foundation for Innovation (CFI), NanoQuébec, RMGA and the Fonds de recherche du Québec - Nature et technologies (FRQ-NT).

## Methods

### Patient data and quality checks

Ethics approval was obtained at the Jewish General Hospital and at McGill University, Montreal, QC, to obtain whole blood samples from the patients with SARDs, at the time of initial diagnosis prior to any treatment. Cell purification and phenotyping Protocols for cell subset isolation, analysis, purity evaluation, fractionation and storage were standardized and optimized. Forty millilitres of peripheral blood were obtained on the above subjects and processed within 4 hours. Peripheral blood mononuclear cells (PBMCs) were separated with Lymphocyte Separation Medium (Mediatech, Inc.). Isolated PBMCs were sequentially incubated with anti-CD19, anti-CD14 and anti-CD4 microbeads (Miltenyi Biotec). Automated cell separation of specific cell subpopulations was performed with auto-MACS using positive selection programs. An aliquot of the specific isolated cell subtypes was used for purity assessment with flow cytometric analysis. A minimum number of 2 million cells from each subpopulations with a purity higher than 95% were frozen in liquid nitrogen for the epigenomic studies. The optimized protocols required the isolation of sufficient numbers of CD4+ lymphocytes (9.04 ± 4.03 × 106) and CD14+ monocytes (7.89±2.96 × 106), and CD19+ B lymphocytes (2.02±1.42 × 106), of sufficient purity to perform the epigenetic analyses. The required number of cells and purity was not always available, especially for the CD19+ B lymphocytes, so we did not have all three cell types for all patients; for this reason the simulation used only two cell types and 46 patients. Illumina Infinium HumanMethylation450 BeadChip data were normalized with funnorm [38]. Also, a number of probes were removed, specifically those on the sex chromosomes as well as probes close to SNPs (ref). There were 375,639 probes remaining after filtering.

Details of the Simulation Method

This simulation design was initially developed in the Masters thesis of the first author [39].

1. **Selection of DMS probes:** *S* = 500 probes were randomly selected to be associated with the phenotype.
2. **Phenotype** (*z_i_, i* = 1, …, n): A random sample of size 46 was drawn from either a Bernouilli distribution (*p* = 0.5) for a binary phenotype, or from a standard normal distribution for a continuous phenotype.
3. **Cell-type-specific methylation values for non DMS probes:** Let *β_ijk_* represent the true methylation value for individual *i*, probe *j* and cell type *k*. For a probe that is not a DMS probe, the simulated value 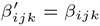.
4. **Cell-type-specific methylation values for DMS probes:**

- Cell-type-specific means are sampled from normal distributions with given parameters. That is, for chosen values *μ_k_*, and *σ_k_*, for *k* = 1, 2, cell-type-specific means for each DMS probe, *μ_jk_*, are generated from 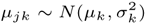.
- The simulated cell-type-specific methylation effect, *ϵ_ijk_*, at a DMS, for an individual sample *i* and an individual probe *j*, is another random quantity, so that

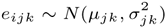
where *σ_jk_* is a parameter provided to the simulation.
- For either binary or continuous phenotype *z_i_*, the simulated methylation value *β_ijk_* is then

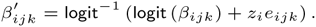 Although all the random effects were simulated on a linear scale, the results are reconverted to the (0,1) scale since several of the cell type adjustment methods require this range.
5. **Combining across cell types:**

- Each individual is assumed to have a unique mixture of the two cell types in a way that depends on the phenotype, *z*. Let 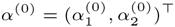 represent the average proportions of the two cell types when *z_i_* = 0, and then let 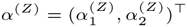 be these proportions when *z_i_* = *Z*. We then say,

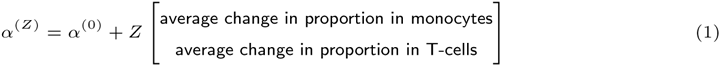
- The cell type proportions *p_ik_* for individual *i* were then generated from Dirichlet 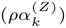, where *ρ* > 0 is a precision parameter, such that larger precision corresponds to less variation in the observed values.
- The final simulated beta value for person *i* at CpG site *j* becomes

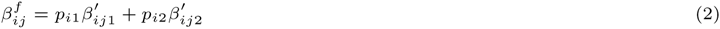

Key notation definitions are summarized in Table 1, and parameter choices for the simulations are in Table 2.

### Description of adjustment methods

Performance of eight popular methods is compared. Brief descriptions of each methods are provided here, and Table 3 compares some key features of the methods, including some details of the implementations. This set of 8 methods is not an exhaustive list of all methods available at this time. In fact, in other fields, particularly engineering and computer science, there exists a plethora of other methods under the guise of “deconvolution” providing the same kind of correction for unmeasured confounding both in other high-throughput data sources [36]. However, we include and compare many of the approaches that are in common usage in the last few years in the world of genomics/epigenomics.

#### Reference-based

This method was published in 2012 by Houseman *et al*. [19]. It relies on the existence of a separate dataset containing methylation measurements on separated cell types. The method uses methylation profiles for the individual cell types to directly estimate the cell type composition of each sample. However, cell-separated data are not always available for all constituent cell types.

#### Reference-free

The second method from Houseman *et al*. does not depend on a reference data set, and therefore can be used in methylation studies on any tissue type [20]. Rather than directly estimating cell type composition, the reference-free method performs a singular value decomposition on the concatenation of the estimated coefficient and residual matrices from an initial, unadjusted model. A set of latent vectors is then obtained that accounts for cell type in further analyses.

#### Surrogate Variable Analysis

Surrogate Variable Analysis (SVA) is a popular method that was introduced by Leek and Storey in 2007 [23]. It was not specifically intended for use in methylation studies, but is nonetheless well-suited for such analyses. SVA seeks a set of surrogate variables that span the same linear space as the unmeasured confounders (i.e. cell type proportions). It is based on a singular value decomposition on the residual matrix from a regression model not accounting for cell type composition. The total number of surrogate variables included in the model is based on a permutation test.

#### Independent Surrogate Variable Analysis (ISVA)

ISVA from Teschendorff *et al*. [24] is very similar in principle to SVA. The main difference is that instead of applying singular value decomposition, it uses Independent Component Analysis (ICA) that attemps to find a set of latent variables that are as statistically independent as possible.

#### FaST-LMM-EWASher (EWASher)

This method from Zou *et al*. [22] extends the Factored Spectrally Transformed Linear Mixed Model algorithm (FaST-LMM) [40] for use in the context of EWAS. A similarity matrix is calculated based on the methylation profiles, and principal components are subsequently included in the linear mixed model until the genomic inflation factor is controlled. The maximum number of principal components allowed was fixed to 10.

#### Removing Unwanted Variation

The method called ”Removing unwanted variation (RUV)” was published in 2012 [26] by Gagnon-Bartsch and Speed. It performs a factor analysis on negative control probes to separate out variation due to unmeasured confounders, while leaving the variation due to the factors of interest intact. Here we use RUV-4, an extension to the original published version, which uses elements from RUV as well as SVA [27]. Control probes were chosen from a list of 500 probes on the 450K platform known to be differentially methylated with blood cell type and age [13]. We selected probes that were not strongly correlated with the simulated phenotype.

#### Deconfounding (Deconf)

The Deconfounding method from Repsilber *et al*. [25] was developed for gene expression studies on heterogeneous tissue samples, but is applicable for use in EWAS. The algorithm performs a non-negative matrix factorization on the methylation matrix, but does not consider the phenotype in correcting for the heterogeneity and does not estimate the number of cell types present.

#### CellCDecon

CellCDecon was developed by Wagner [21], and is similar to Deconfounding in that it does not consider the phenotype in performing its decomposition and does not internally estimate the number of cell types present. The method assumes a specific regression parameterization, and makes random perturbations to the model parameters which are accepted if there is a decrease in the sum squared residuals.

**Supplement Figure 1:**
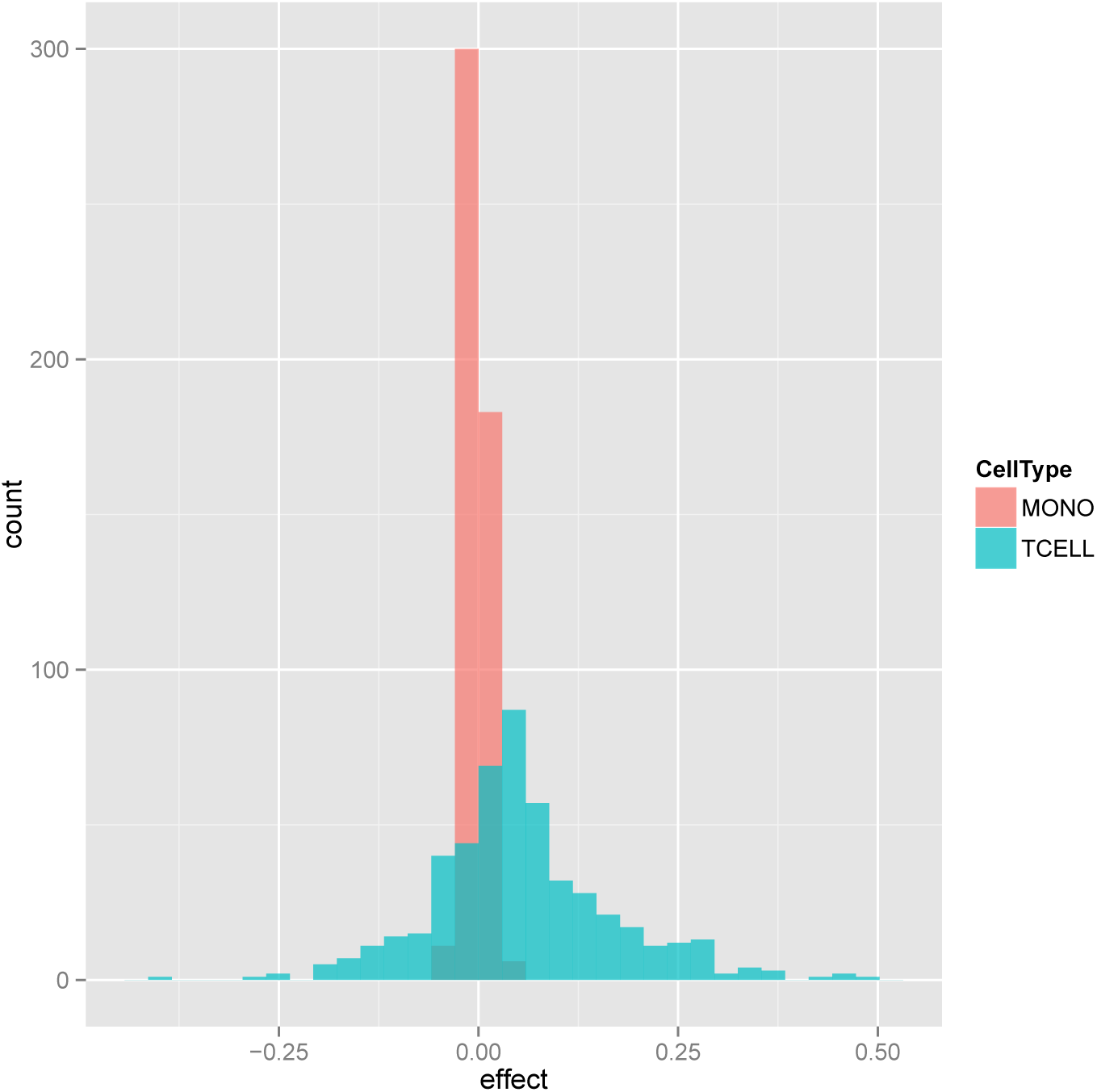
Histograms of simulated effect means *μ_jk_* for truly associated probes in Simulation Scenario 1

**Supplement Figure 2:**
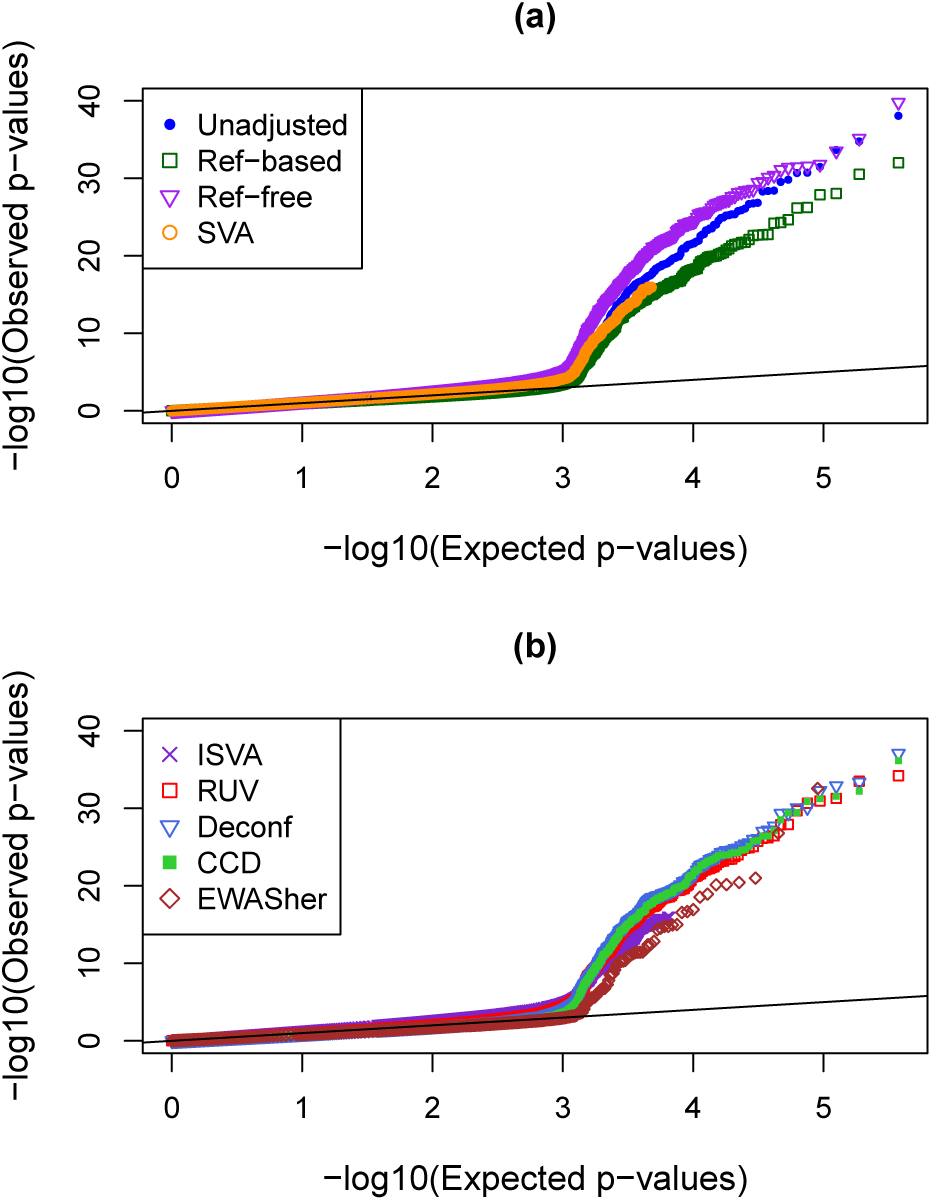
QQ-plots showing distributions of p-values in when there is no confounding (Simulation Scenario 2).

**Supplement Figure 3:**
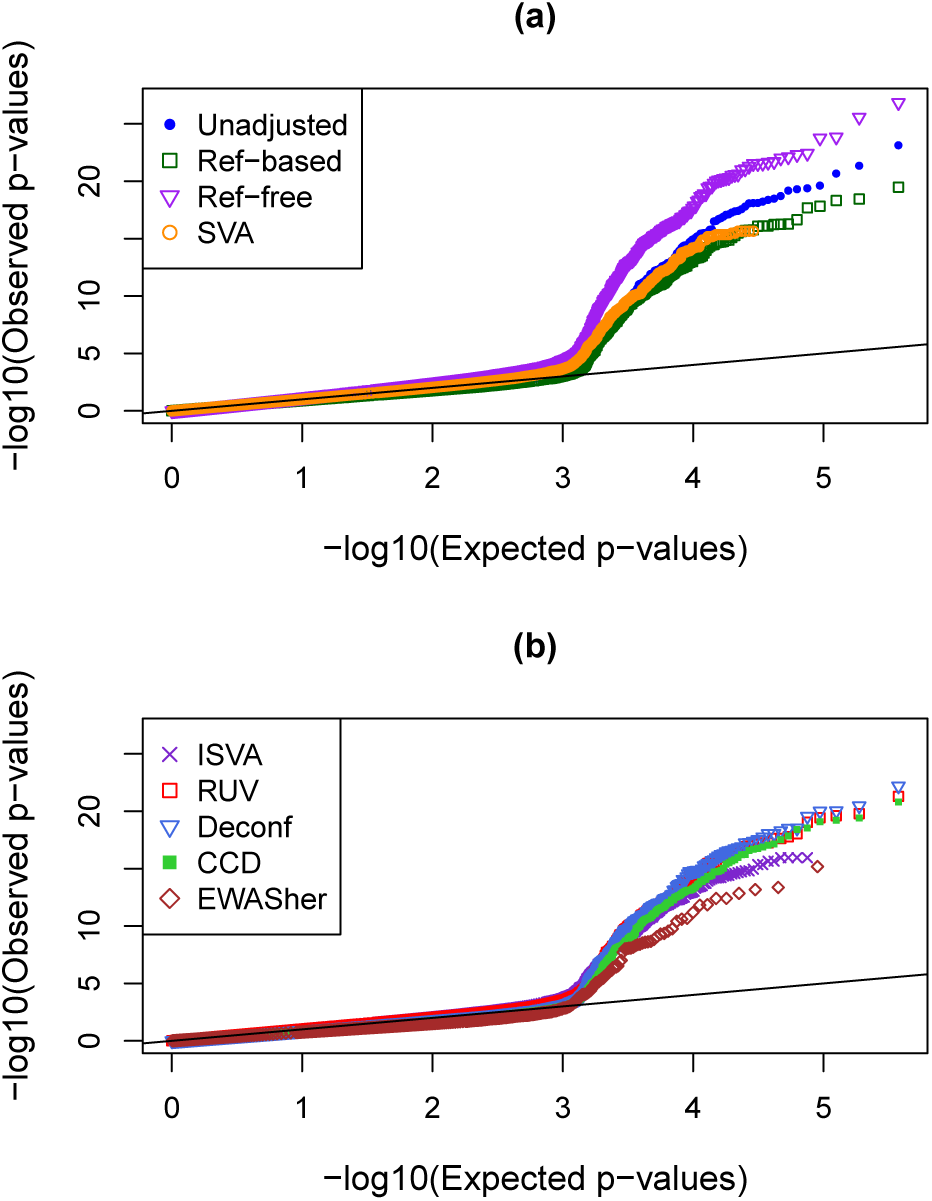
QQ-plots showing distributions of p-values in when there are opposite effects (Simulation Scenario 3).

**Supplement Figure 4:**
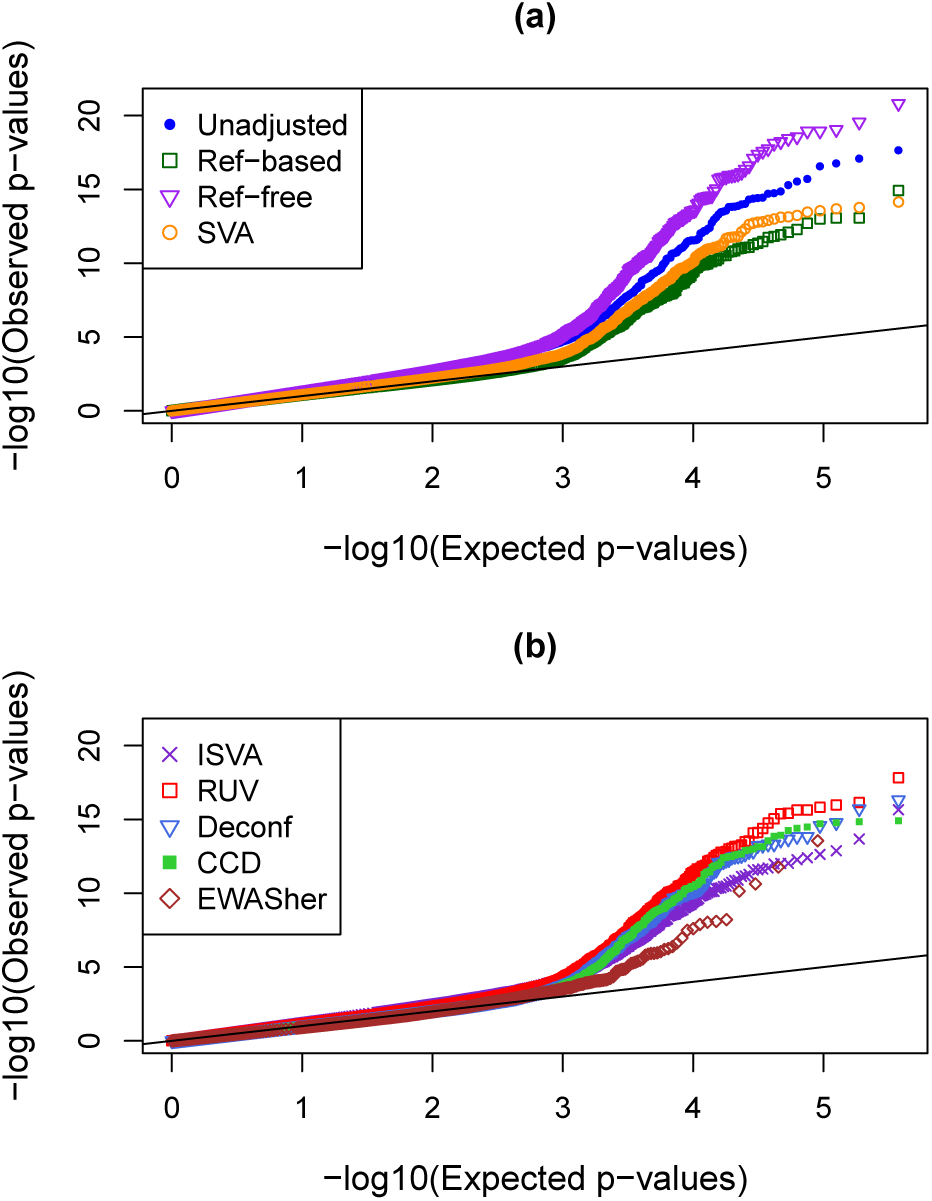
QQ-plots showing distributions of p-values in when there is high precision (Simulation Scenario 4).

**Supplement Figure 5:**
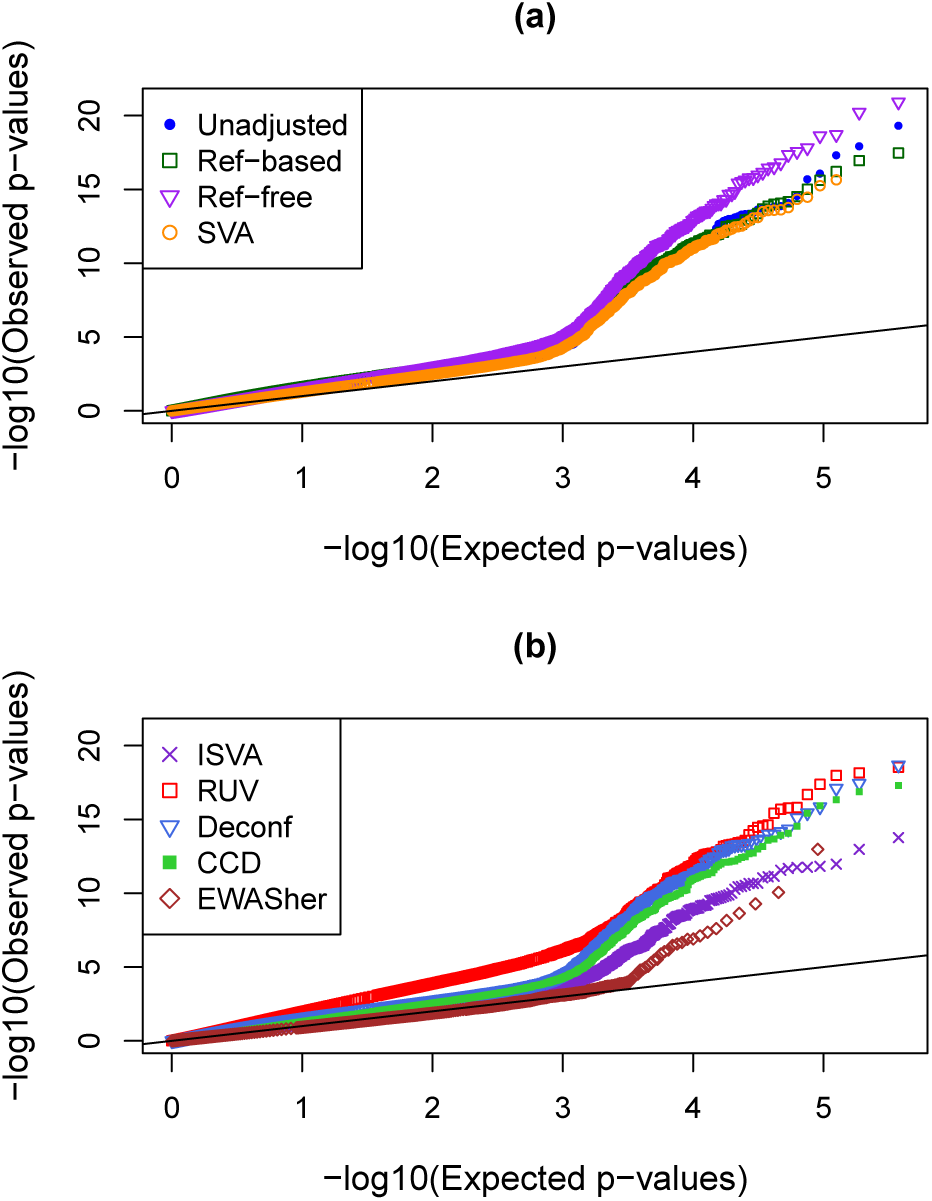
QQ-plots showing distributions of p-values in when there is low precision (Simulation Scenario 5).

**Supplement Figure 6:**
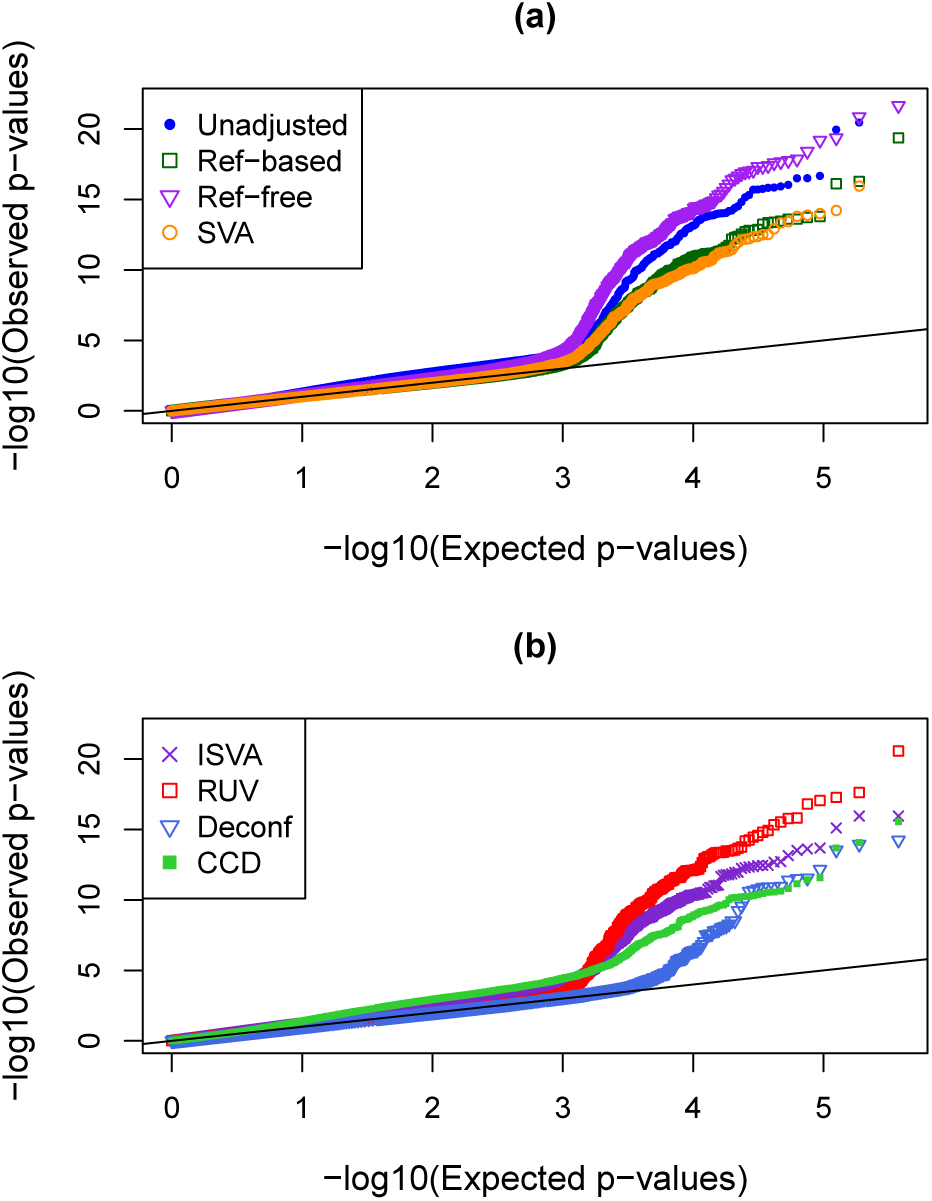
QQ-plots showing distributions of p-values for continuous phenotypes (Simulation Scenario 6).

